# Temporal interference electrical neurostimulation yields fMRI BOLD activation in humans

**DOI:** 10.1101/2024.03.02.583125

**Authors:** Priyamvada Modak, Justin Fine, Brayden Colon, Ella Need, Leslie Hulvershorn, Peter Finn, Joshua W Brown

## Abstract

Temporal interference electrical neurostimulation (TI) is a relatively new method of non-invasive neurostimulation that may be able to stimulate deep brain regions without stimulating the overlying superficial regions. Despite studies in rodents, almost no studies have investigated its effects on human brain activity along with safety and tolerability profiles. We performed simultaneous TI stimulation and fMRI to investigate the effects of TI on human BOLD signals. Here we show that TI can induce increased BOLD activation in humans, with good safety and tolerability profiles. We also show the limits of spatial precision and explore the nature and causes of additional off target effects. TI may be a promising approach for addressing questions about the causal role of deep brain structures in human cognition and may also afford new clinical treatments.

## 1. Introduction

The primary means to selectively stimulate deep brain regions, until very recently, was to invasively implant electrodes in the brain. In 2017, a new technique, temporal interference electrical neurostimulation (TI), was proposed by Grossman et al. (2017) that showed promise for non-invasively targeting deep brain regions. TI involves two high frequency alternating currents with slightly different frequencies, typically > 2Khz. These currents presumably do not cause significant neurostimulation individually, however, they produce an interference pattern whose low frequency component apparently causes neurostimulation at the location where the interference occurs. TI holds great promise for the field of cognitive neuroscience because apart from clinical applications, the ability to non-invasively stimulate deep brain regions can be useful in determining the causal role of deeper brain regions in various cognitive functions.

Grossman et al. (2017) tested and validated the technique in mice. Acerbo et al. (2022) then showed that high-frequency TI affects sharp-wave ripples in the hippocampus and suppresses epileptic biomarkers in mouse models. Esmaeilpour et al. (2021) showed that TI can modulate gamma oscillations in rat hippocampal slices, that GABA-mediated adaption mechanisms are central to the TI response, and how the specific selection of carrier frequency may affect the sensitivity and selectivity of TI. Despite these investigations in animals, it remains unclear whether TI had any off-target effects.

There have been some previous studies which have used TI in humans (Ma et al., 2022; Zhu et al., 2022). These studies show that the TI is effective in bringing about an observable change in behavior, target region excitability and functional connectivity of the target region, however, it remains to be determined how effective is TI in stimulating the target region and how selective or focal that effect is. Further, Acerbo et al. (2022) showed that TI could be delivered to humans in the hippocampus, as demonstrated by recording the generated electric fields inside human cadavers using stereo-electrode-encephalography electrodes.

To date, only one study (Violante et al., 2023) has validated the effect of TI (targeted at hippocampus) on neural activity and memory in humans, and, overall, data on the safety and tolerability of TI in humans is sparse.

We targeted the left Caudate/Nucleus Accumbens (NAc) in this study because, if validated, we hope to stimulate the NAc using TI to explore its role in substance use disorders and in specifically disrupting substance use behavior. Previous research has shown that lesions or invasive deep brain stimulation of NAc has led to improvements in relief from drug seeking behavior (Müller et al., 2009a, 2009b). Therefore, an ability to non-invasively target NAc would open avenues to easily investigate its causal role in substance abuse as well as develop treatments.

Thus, the objective with this study was to explore whether TI in humans can cause changes in the fMRI BOLD signal, and how focal (selective) the stimulation is, and identify any potential adverse effects along with general safety and tolerability. Specifically, we hypothesized that TI neurostimulation of NAc would lead to BOLD activation. Secondarily, we sought to characterize how focal the effects are, and how spatially variable they are in this region. Further, what is the (potentially nonlinear) relationship between the TI electrical field and the BOLD signal? The analysis of relationship between TI and BOLD signal was exploratory as there is no prior research that would support a specific hypothesis about the nature of this relation.

## 2. Materials and methods

### 2.1 Participants and study procedures

A total of 16 healthy subjects (Age: mean = 28.9 years, standard deviation = 10; Sex: 10 females and 6 males) participated in the study. Two pairs of MRI-compatible carbon electrodes placed at {F9,F10} and {Fp1, CPz} were used to administer neurostimulation. Subjects completed measures of mood (Watson & Clark, 1999), working memory (Digit span; Richardson (2007)), and a newly developed Stimulation Experience Scale (SES) throughout the neurostimulation and fMRI session. Please see Supplementary sections F.1 and F.3 for more details.

### 2.2 Neurostimulation specifications and finite element analysis

The neurostimulation apparatus specified in section F.2 in the Supplementary materials was used to administer high frequency alternating electric current to the scalp of the subjects.

The electrode locations were optimized for one subject which we refer to as the **reference subject** from here on. This was done using the SimNIBS software (Saturnino et al., 2019) to create a finite element model (FEM) of the brain from the structural images of the subject. The optimization function maximized a measure TI_fm_ (See Supplementary F.6) such that the TI magnitude in target region was maximized while minimizing it in all other regions. The target and non-target regions were defined using the neuromorphometrics atlas, Neuromorphometrics, Inc., Winthrop, MA, USA.

All subjects were stimulated using the optimal electrode locations found above, however, we also performed finite element analysis for each subject using their structural images to find predicted TI in each subject brain. We created an image of the mean minus two standard deviations of the predicted TI values across the subjects for every voxel in MNI space. We refer to this image as the ‘**all-subjects TI**’ image, representing the lower estimate of the predicted TI effect at a particular voxel, and we will refer to its voxel values as TI^*^ in the rest of the paper. The TI* was high (>0.6) in the frontal lobe, so we created a mask with MNI y>22 which we used to perform small volume correction of the fMRI results. The details of the electrode optimization, rationale for the all-subjects TI image, and the above mask can be found in the Supplementary material (F.6, F.7, and F.8).

### 2.3 Experiment design

We conducted four 8-min blocks with alternating blocks of <u>active</u> and <u>sham</u> stimulation such that there were two blocks of active stimulation and two blocks of sham stimulation. The active stimulation block consisted of 120 seconds of stimulation (which we refer to as ‘ON’) alternating with 120 seconds of no stimulation where zero current was passed through the electrodes (which we refer to as ‘OFF’). The ON period consisted of 30 seconds of current ramp up from 0 to 2 mA. The current ramped down to 0 mA in the first 30 seconds of the OFF period.

The sham block had alternations between ON and OFF just like the active block, however, the ON in the sham did not have the 60 seconds of stimulation between ramp-up and ramp-down. It only consisted of 30 seconds of ramping up from 0 to 2 mA followed by immediate ramp down from 2 to 0 mA. The sham block controlled for the sensations on the scalp that the subjects feel/may feel when being stimulated, which may occur during the ramp up (Gandiga et al., 2006).

A particular block could either begin with an ‘ON’ or an ‘OFF’ period, and we refer to blocks as ON-block or an OFF-block depending on what they began with. The active ON-block and its corresponding sham ON-block (figure 1 A, B) were conducted consecutively, in any order. The same was done in the active OFF-block, and its corresponding sham OFF-block. Further, no two sham blocks or two active blocks, irrespective of whether they were ON-or OFF-blocks, could be together.

**Figure 1:**
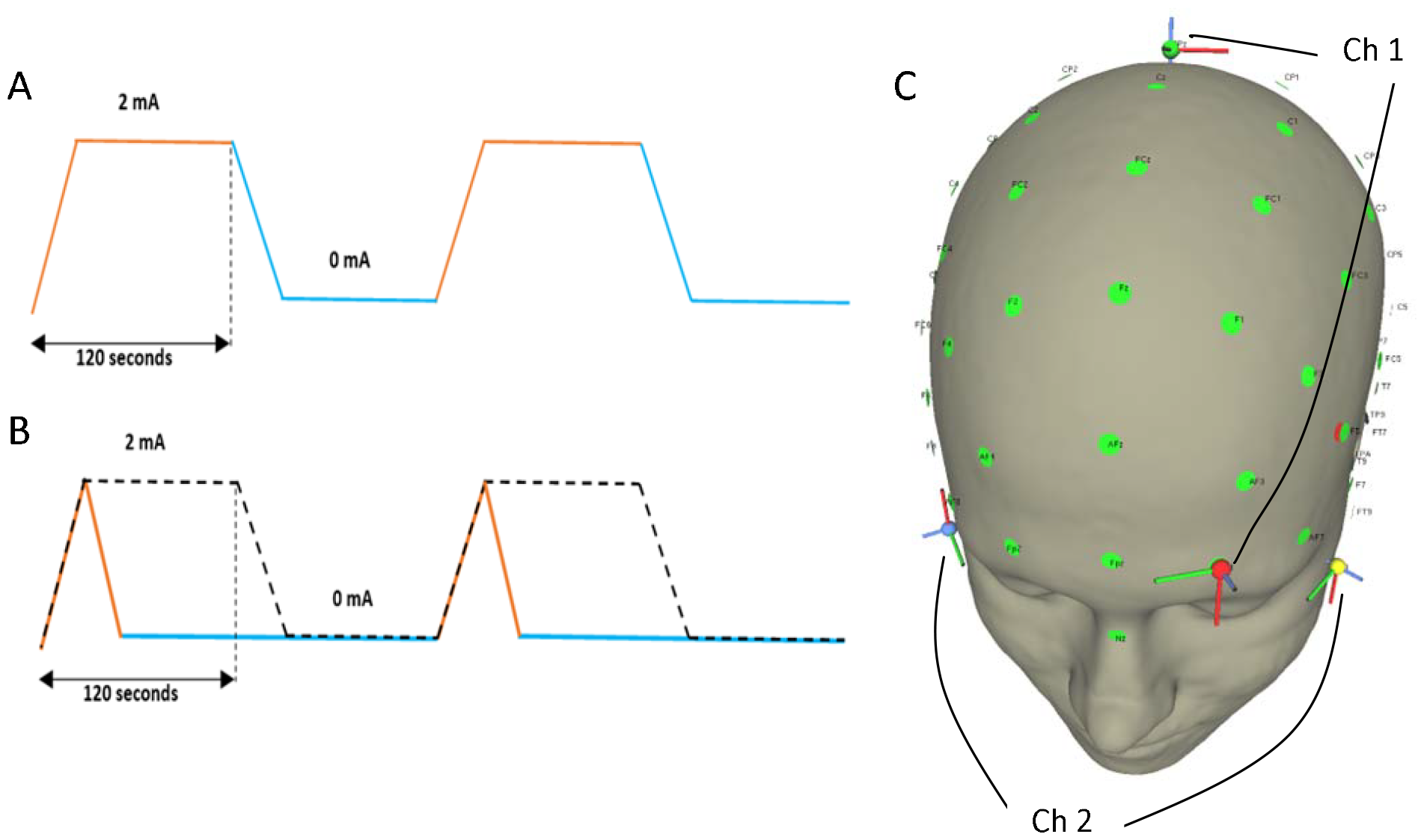
A - Visual depiction of an ON-active-block - that is, an active-block (either TI or no-TI) that began with the ‘ON’ stimulation. Figure shows the changes in current passed through a pair of electrodes throughout the block. Here, orange - ‘ON’ (120 seconds); Blue - ‘OFF’ (120 seconds); B - Visual depiction of an ON-sham-block - that is, a sham-block (either TI or no-TI) that began with the ‘ON’ stimulation. Orange and blue lines show the changes in current passed through a pair of electrodes throughout the block. The dotted black line shows the sham regressor as considered in the GLM; C – Electrode locations of stimulation (F9 and F10, FP1 and CPz).

Further a block could either be a TI-block or a no-TI-block. In the TI condition, the currents through two pairs of electrodes were administered at frequencies 2000 Hz and 2020 Hz so that the interference pattern of their induced electric fields was expected to have a low frequency component of 20 Hz. In the no-TI-condition, both the currents were administered at 2000 Hz, so that their interference pattern would have no low frequency component, that is, there was ‘no TI’. The no-TI-condition controlled for any effects of high frequency currents on the fMRI BOLD activation (Mirzakhalili et al., 2020) thus allowing us to identify BOLD effects of TI as distinct from BOLD effects that may be due to the individual high frequency stimulation. Please refer to section F of the supplementary material for more details.

### 2.4 fMRI analysis: General linear model

The fMRI acquisition and preprocessing methods have been described previously (Modak et al., 2021). We analyzed the fMRI data using SPM12 (supplementary material F.5). We formed the following regressors: ON_Active_TI, ON_Active_No_TI, ON_Sham_TI, ON_Sham_No_TI, OFF_Active_TI, OFF_Active_No_TI, OFF_Sham_TI, and OFF_Sham_No_TI. These regressors each spanned the entire duration of a block, and the magnitude of the regressor at each point in time was simply the electrical current magnitude, which was the same for each electrode pair (Figure 1 A, B), convolved with a canonical hemodynamic response function. For the sham condition, we modeled the time course exactly as if it had been the active condition rather than following the actual current in the sham condition. We reasoned that this allows a direct comparison of the BOLD signal for same time points with vs. without stimulation.

We constructed the following contrasts of interest:

C1: BOLD beta weight contrast of (StimTI – ShamTI). Here, StimTI consisted of regressors ON_Active_TI, and OFF_Active_TI. ShamTI consisted of regressors ON_Sham_TI, and OFF_Sham_TI.

C2: BOLD beta weight contrast of (StimTI – ShamTI) – (StimNoTI – ShamNoTI). Here, StimNoTI consisted of regressors ON_Active_No_TI, and OFF_Active_No_TI. ShamNoTI consisted of regressors ON_Sham_No_TI and OFF_Sham_No_TI.

## 3. Results

### 3.1 Safety and tolerability

Across the 16 subjects, there were a total of 23 blocks of stim-TI and sham-TI, each. The last 8 subjects also were administered the no-TI condition and hence they only had one block each of stim-TI and sham-TI (Table S4). The average total score on the SES scale for the stim-TI blocks was 12.04 and for the sham-TI blocks was 12.35 where the total score could range from 11 to 55 where 11 indicates no discomfort and 55 indicates maximum discomfort. The ‘discomfort’ here was assessed in terms of ‘stress’, ‘anxiety’, ‘ringing in ears’, ‘headache’, ‘itchiness’, etc. The raw self-report scores on each of these are provided in the supplementary material (Table S4). The total scores in stim-TI and sham-TI blocks did not differ significantly (p-value = 0.58; Wilcoxon-signed ranked test). Table S5 also shows the number of stim-TI and sham-TI blocks for each of the categories of the SES scale for which some discomfort was reported.

All but one of the subjects reported no long-term effects of the stimulation and scanning session in the follow-up call made 3-15 days (mean = 9 days) after the participation. One subject reported experiencing slight headaches post 15 days of the session.

### 3.2 BOLD fMRI activation effects

We looked at the effect of StimTI – ShamTI contrast. The effect of this contrast is expected to reveal the BOLD activation due to TI as well as due to any effect of the high frequency electric currents.

We see two clusters showing a strong positive effect on this contrast. One in bilateral mid-orbito-frontal cortex (OFC) (figure 2A), and one in the fusiform/parahippocampal region (figure 3). The latter is significant after whole brain cluster correction while the former is not but is close to being significant at cluster corrected p-value of 0.092. This cluster, however, is significant after small volume correction using the frontal mask shown in figure 2B.

**Figure 2:**
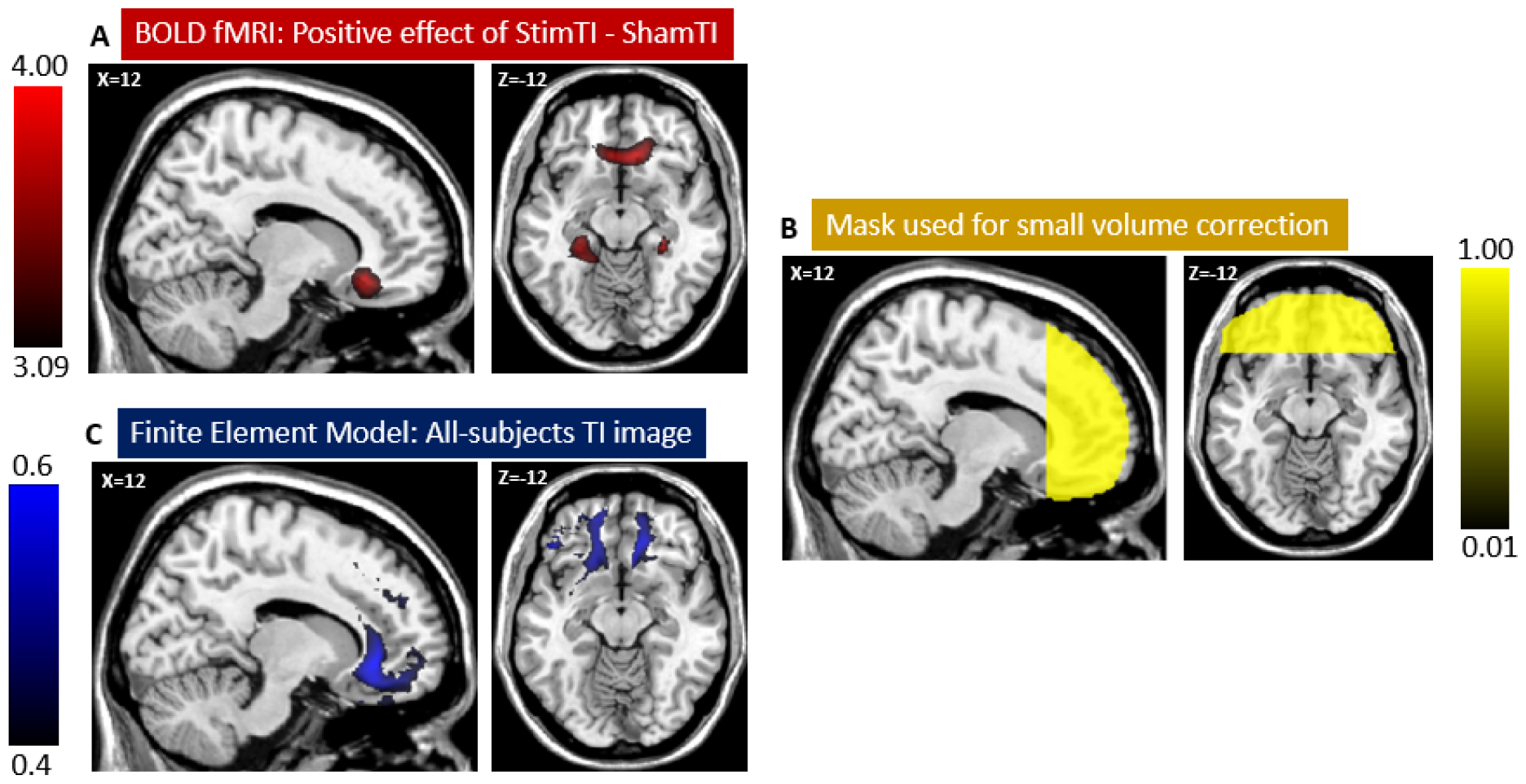
**A – Positive effect of contrast StimTI – ShamTI** in OFC; B – Mask used for small volume correction; C – all-subjects TI image

**Figure 3:**
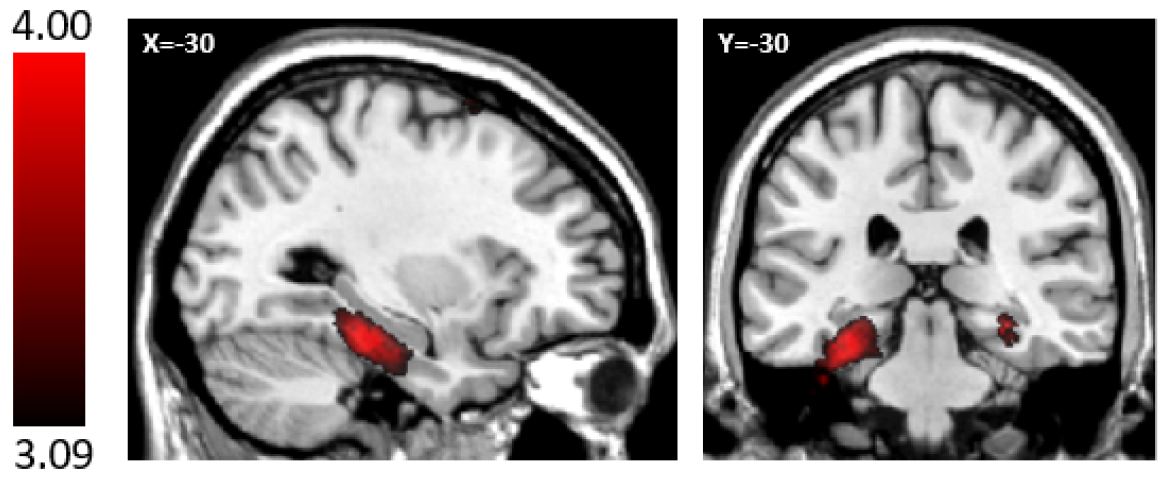
**Positive effect of contrast StimTI – ShamTI** in parahippocampal gyrus

### 3.3 BOLD fMRI suppression effects

We examined the possibility that the high frequency current alone might reduce neural activity, perhaps by blocking conduction (Mirzakhalili et al., 2020). To explore this, we calculated the negative effects of the StimTI – ShamTI contrast, which showed a weak effect of suppression in the right precuneus and superior parietal lobule. These were not significant at the whole brain cluster correction level. Nevertheless, we noted that this region roughly coincided with the location of the CPz electrode, so we explored further to ascertain whether the cluster might be significant with a small volume correction around the CPz electrode.

As shown in table 3, the parietal cluster is significant at p = 0.036 after small volume correction using the mask of radius 30 mm around the average location of CPz electrode across the subjects as shown in right panel of figure 4. Please refer to section F.8 in supplementary material for details on this mask. This finding should be interpreted with caution given that it was only significant within a small volume around the electrode, and similar effects were not found around the other three electrodes, even with similar masks.

**Table 1:**
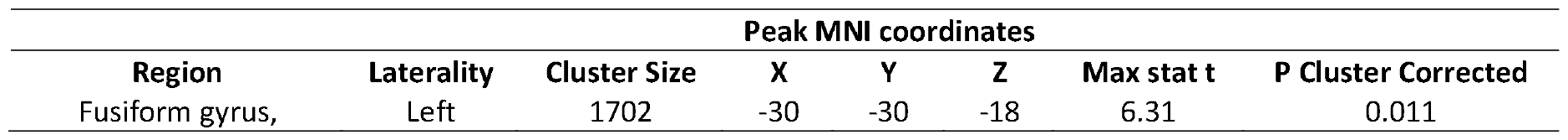
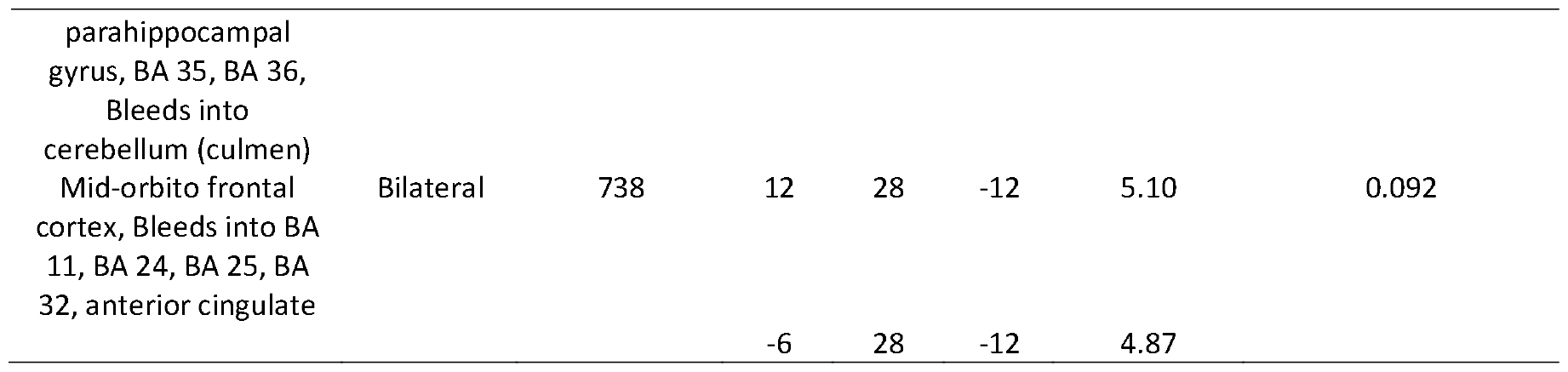
Positive effects of StimTI – ShamTI contrast. Whole brain cluster correction. Cluster-defining threshold = 0.001.

**Table 2:** Positive effects of StimTI – ShamTI contrast. Small volume correction using the frontal mask, y > 22. Cluster-defining threshold = 0.001.

| Region | Laterality | Cluster Size | Peak MNI coordinates |  |  | Max stat t | P Cluster Corrected |
| --- | --- | --- | --- | --- | --- | --- | --- |
|  |  |  | X | Y | Z |  |  |
| Mid-orbito frontal cortex, Bleeds into BA 11, BA 24, BA 25, BA 32, anterior cingulate | Bilateral | 712 | 12 | 28 | -12 | 5.10 | 0.021 |
|  |  |  | -6 | 28 | -12 | 4.87 |  |

**Table 3:** Negative effects of StimTI – ShamTI contrast. Small volume correction using the mask around the location of Cpz electrode. Cluster-defining threshold = 0.001; Cluster extent threshold = 5.

| Region | Laterality | Cluster Size | Peak MNI coordinates |  |  | Max stat t | P Cluster Corrected |
| --- | --- | --- | --- | --- | --- | --- | --- |
|  |  |  | X | Y | Z |  |  |
| Precuneus, Superior parietal lobule, BA 7 | Right | 110 | 8 | -76 | 64 | 3.97 | 0.036 |

**Figure 4:**
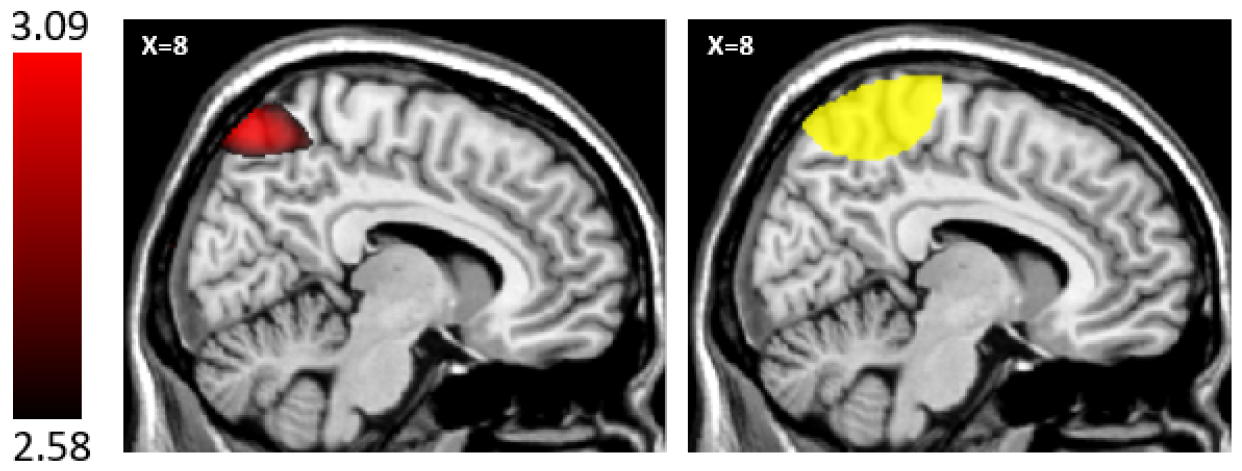
Negative effects of StimTI – ShamTI contrast. Left – Region in right precuneus and superior parietal lobule which is significant after small volume correction using the mask on the right. Right – Mask used for small volume correction – it is an overlap of the sphere of radius 30 mm around [0, -52, 78] and the brain (that is, the mask used for StimTI-ShamTI second-level analysis).

### 3.4 Spatial localization of BOLD TI effects

The region that shows BOLD activation in the frontal cortex varies slightly from the one that we intended to target (Figure S2). This raises the question about efficacy and spatial localization of TI. If the TI field is high at a certain location, does that necessarily mean that there will be BOLD signal change at that location in the brain?

It should be noted that we performed the optimization of the stimulation set-up for a reference subject, so it is possible that a different region was targeted when it was applied to other subjects. To assess this, we created a head model and performed the finite element analysis individually for each subject using which we constructed an all-subjects TI image, as explained in section 2.2.

As can be seen in figures 2 and S2, the region with high values of TI^*^ is adjacent to the region that we intended to target but not perfectly overlapping. We created a mask of the frontal region containing these high TI^*^ values (Figure 2B) and used it to perform small volume correction of the BOLD activation observed in frontal region. The results showed a region in frontal cortex with significant BOLD activation which overlapped with the region showing high TI^*^ values (Figures 2A,C). The threshold chosen to define ‘high’ in the above should ideally have been the minimum TI value necessary to induce BOLD activation, but since we don’t know the functional relationship between TI field strength and BOLD activation (see Supplementary material section C), we arbitrarily chose TI^*^ >= 0.6 V/m to define the region with ‘high’ predicted TI effects (section 2.2 and Supplementary F.8). This threshold was chosen before we performed the small volume correction mentioned above. In mice, Grossman et al. (2017) showed that TI can cause stimulation for scalp currents as low as 100 microamperes while in rats. Esmaeilpour et al. (2021) explore the electric fields necessary to bring about changes in gamma power. It seems from our study that values as low as 0.28 V/m (TI^*^= 0.08 V/m) may be able to lead to BOLD activation, however, it remains to be determined whether these stimulations were direct or indirect effects of TI. The value 0.28 V/m is the average value of TI across the subjects in the parahippocampal region showing significant BOLD activation. Supplementary figure S2 shows the overlap between the high TI^*^ region and the cluster in frontal cortex mentioned above.

The mean value of TI^*^ in the cluster in the frontal cortex that is significant after small volume correction is 0.32 V/m. This value for the target region (defined using neuromorphometrics atlas) is 0.22 V/m which is smaller than the one in the above cluster.

Even in the reference subject, neither the TI_fm_ nor the mean TI^*^ is highest in the target region vs. other regions. This is because the optimization for the reference subject was not for the mean TI in the target region but for the TI_fm_ mentioned in section 2.2, and the optimization was performed to select from candidate electrode positions that had the highest TI_fm_ possible in the left-caudate/NAc but not necessarily the highest TI_fm_ in the target region among all the regions. TI_fm_ is the highest in the left medial orbital gyrus in the reference subject. The mean TI^*^ values for atlas-defined regions in the reference subject also show a similar pattern as in the all-subjects TI image above (Figure 2C). The TI prediction for the reference subject is not all that different from the TI predictions for the study participants, which suggests that it may suffice to optimize stimulation set-up for a reference subject like we have done here. We explore this further in section 3.6.

So, we find that where the TI is high, there is BOLD activation suggesting that TI is efficacious in inducing BOLD activation, at least with the 20Hz frequency difference used here. This is in line with the results of Grossman et al. (2017) who showed the efficacy of TI in mice. However, it does not necessarily mean the converse, i.e. that wherever there is BOLD activation, the TI is high as we also see BOLD activation in parahippocampal gyrus, where the TI is relatively low. The significant cluster in parahippocampal gyrus has a mean TI^*^ of 0.08 V/m which is smaller than the mean TI^*^ in the original target region which did not show a significant BOLD activation.

### 3.5 What is the functional relationship between TI and BOLD?

We found above that there is BOLD activation where the TI* is relatively high, but the exact functional relationship between them is less clear (details in Supplementary material C). The answer to this question might allow future improvements to the algorithms used to optimize electrode placement and TI focality within the brain. We considered all gray matter voxels and asked what functional form provides the best account of how increasing TI leads to increasing BOLD signals. We find that the BOLD activation for stimTI – shamTI was best predicted by a logistic function of the TI values predicted from finite element modeling at the level of voxels. One challenge in finding a model of TI that would reliably vary with BOLD activation across subjects was that not all subjects had a significant positive correlation between TI and BOLD.

### 3.6 Is it necessary to optimize electrode placement for each individual?

We next explored whether it is beneficial to optimize the stimulation setup individually for every subject rather than simply for one reference subject.

The factors that affect the TI field are head shape and volume, tissue conductivities, and tissue segmentation. To target a particular region in a subject, one should ideally collect the structural MRI and diffusion MRI of that subject and perform the FEM analysis using these, including anisotropies. If the EEG cap is being used for electrode placement, then one could additionally incorporate the shift in position of electrodes due to the stretching of the cap. However, how worthwhile it is to undertake this effort?

Even though we did not control for these sources of variation, the predicted TI values of subjects are highly correlated with each other (Supplementary material B). For example, the correlation across voxels between the predicted TI values in the reference subject and average of the predicted TI values across the subjects is 0.6043 (p<0.001). To further address this question, we looked at the variance across the subjects in the peak MNI coordinates of their predicted TI value (via FEM) as well as that of the stimTI – shamTI contrast (with a 20mm smoothing kernel).

For every subject, we created a sphere of radius 10 mm around the peak MNI coordinates of the predicted TI. The blue in the right panel of Figure 5 shows an overlap of these spheres for various subjects. It has the highest value (16) for voxels that are part of all the spheres while lowest (1) for voxels that are part of only one sphere. The same was done for the peak MNI coordinates of the (stimTI – shamTI) BOLD contrast but after masking the contrast images with the frontal mask (Figure 2B). See supplementary figure S8 to view the peak MNI coordinates without this masking. The reference subject was apparently an outlier in that the MNI Z-coordinate value was lower than that of the other subjects.

**Figure 5:**
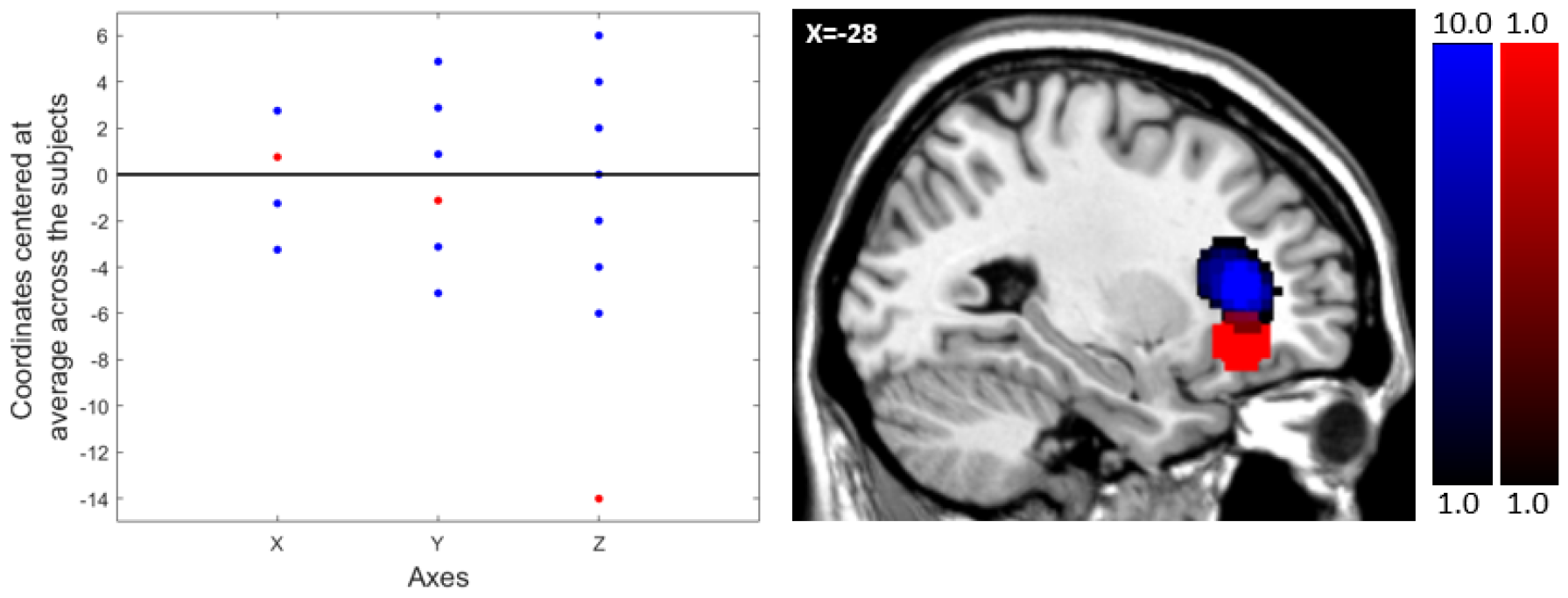
Variability in peak MNI coordinates for predicted TI across subjects. Left: Plot shows the peak MNI coordinates of the predicted TI for subjects. The X, Y, Z coordinates are separately centered at their means across the subjects and are shown as blue dots. The red dots correspond to the peak MNI coordinates of predicted TI in the reference subject which is centered using the above mean. Right: Blue shows the overlap of 10 mm spheres around the peak MNI coordinates of the subjects while red shows the 10 mm sphere around the peak MNI coordinates of the reference subject.

The standard deviation in peak MNI coordinates for X, Y, and Z coordinates for predicted TI were 1.77, 3.34, and 3.18 mm (Figure 5) and for BOLD contrast were 36.87, 13.20, and 20.81 mm (Figure 6), which suggests that the variability across subjects in the predicted TI peak is low relative to the variability in the observed fMRI BOLD peak.

**Figure 6:**
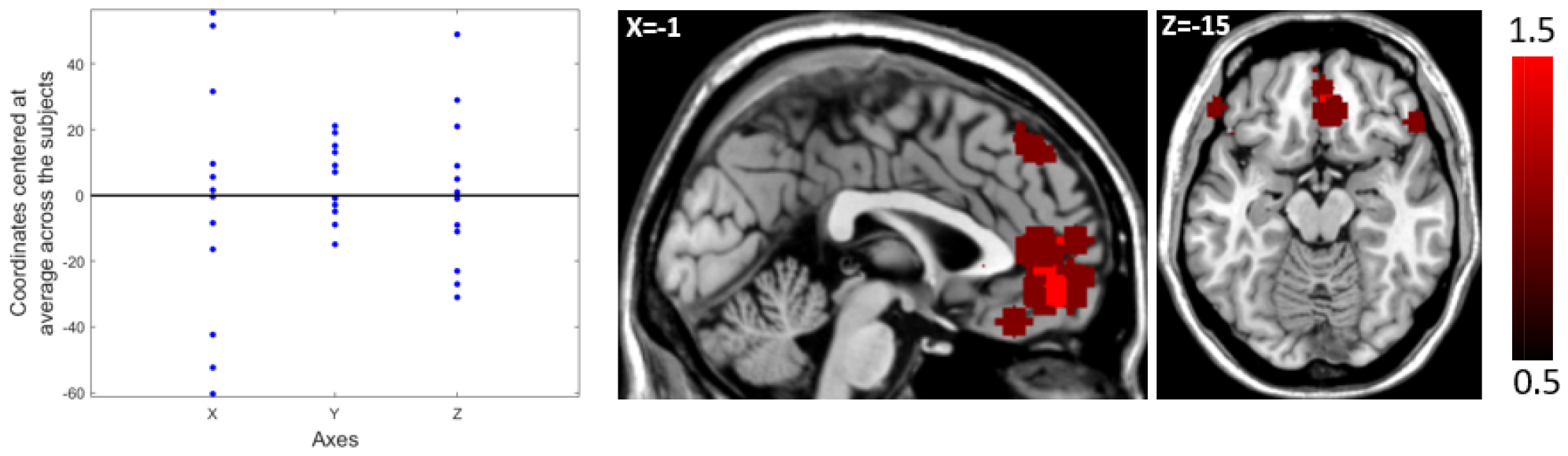
**Variability in peak MNI coordinates for BOLD contrast values for (stimTI – shamTI) across subjects** and masked by the frontal mask (figure 2B). Left: Plot shows the peak MNI coordinates of the BOLD contrast for subjects. The X, Y, Z coordinates are separately centered at their means across the subjects and are shown as blue dots. Right: Red shows the overlap of 10 mm spheres around the peak MNI coordinates of the subjects.

### 3.7 Potential cognitive confounds on BOLD

We also considered the possibility that the observed effects of stimTI – shamTI contrast could be a result of cognitive effects of stimulation procedure apart from the stimulation itself such as the effect of the feeling of getting stimulated. However, our results from analyzing subjects’ ratings from stimulation experience scale (SES) did not show any significant differences between stimulation and sham conditions (section 3.1). We cannot rule out, however, that there were differences in experience of both conditions which were not captured by the SES.

## 4 Discussion

In this work, we set out to test whether the temporal interference neurostimulation (TI) proposed by Grossman et al. (2017) would modulate the fMRI BOLD signals in humans. We targeted the left Caudate/Nucleus accumbens (NAc) using TI while simultaneously collecting BOLD fMRI data in healthy subjects. We found that regions in the bilateral orbitofrontal cortex (OFC), which is close to the NAc, showed BOLD activation as the main effect in the targeted regions. We also found that TI may result in accompanying activation or deactivation in off-target regions. We also collected the subjects’ self-reports on the new stimulation experience scale (SES) while they were being stimulated to assess the safety and tolerability of the TI-stimulation. Our results indicate that TI-stimulation leads to various BOLD effects and does not pose a significant safety risk to the subjects.

What accounts for the BOLD effects in targeted regions (regions with high TI)? Cao & Grover (2020) simulated the biophysics of TI effects on neurons and show that increasing TI signals may cause sodium channels to open. This does not necessarily lead to net activation of the tissue, as that depends on whether the activated neurons are predominantly excitatory or inhibitory.

Grossman et al. (2017b) found evidence of increased metabolic activity in TI stimulated regions in rodents, but more recently Violante et al. (2023) found that TI with a 5 Hz beat frequency in the theta band (vs. the 20Hz beat frequency in the beta band here) led to reduced BOLD signal in the hippocampal target region vs. the increased BOLD signal we found here. It may be that different beat frequencies can produce excitation vs. inhibition.

There are at least four possible accounts of off-target effects, namely pure alternating current effects, functional connectivity effects, tissue response function differences, and cognitive effects. Regarding pure alternating current effects, Mirzakhalili et al. (2020) show, through simulations of Hodgkin Huxley model of neuron, that the high frequency currents could lead to conduction blocks in the axon. This may explain the deactivation in the region in precuneus/superior parietal lobule which was within 30 mm around the average location of the Cpz electrode (Figure 6). As this region is close to the Cpz electrode, a conduction block is a plausible cause of the deactivation (Mirzakhalili et al. (2020)). The caveat here is that we did not find a region of deactivation within 30 mm of the remaining three electrodes. However, there was also more predicted TI in the vicinity of other electrodes than Cpz which may have countered the deactivating effect of the high frequency currents (Supplementary Figure S6).

Pure alternating current effects are unlikely to account for BOLD activation in regions farther from the electrodes, such as we observed in the parahippocampal region. Mirzakhalili et al. (2020) present a “sandwich” hypothesis wherein some neurons that are off target and have a low (but not zero) magnitude of TI would show tonic activity as opposed to the phasic activity in neurons in the target region with a high magnitude of TI. The possibility of such a tonic activity in parahippocampal region is another candidate reason for observing its BOLD activation.

Alternatively, functional connectivity between the TI target region in the OFC and the parahippocampal region may account for the increased parahippocampal BOLD signal. This remains a question for future work. Further, it is interesting to note that in our study as well as in other studies of TI (Acerbo et al., 2022; Grossman et al., 2017c), activation in temporal lobe was found. Ma et al. (2022) and Zhu et al. (2022) targeted the motor cortex, but it is not known what other regions may have been activated. In Acerbo et al. (2022) and Grossman et al. (2017b), the hippocampus was indeed the target in contrast to our study.

Further different regions may have different excitability thresholds making them susceptible to BOLD activation even at low TI values. Excitability may be governed by proportions of excitatory and inhibitory neurons in a region (Esmaeilpour et al., 2021).

Another reason could be confounding factors in the experiment itself, although we consider this unlikely. In particular, if the subjects experienced the active and sham stimulation differently, the different perceptual experiences may have led to differential BOLD activities. We considered that the itching and tingling on the scalp may be most pronounced during the first few seconds of the stimulation (Gandiga et al., 2006), which is why we briefly activated the current during the start of the sham stimulation, so that the perceptual experiences would match as closely as possible. We found that the SES measures do not differ significantly between active TI and sham TI conditions suggesting that the experience of neurostimulation in both cases was not significantly different.

Nevertheless, could an interaction of experience and the GLM regressors somehow have led to different BOLD signals? In the sham condition, the current ramps down while the GLM regressor is high, but in the active condition, the current ramps down as the GLM regressor also ramps down (Figure 1 A, B). If the current ramp-down in the sham condition caused a stronger sensation, that could in principle lead to BOLD signals reflecting that in the sham condition. To address this, we further compared the BOLD activations of blocks regressors without parametrically modulating them, which avoids minimizing the current ramp-down effects in the active condition. We did not find any significant effects of this contrast.

In a subset of subjects, we implemented a No-TI condition which only has high frequency currents and not the low-frequency component in TI. If the activation in parahippocampal gyrus is an effect of high frequency currents, then we expect it to show positive effect of a contrast such as stimNoTI – shamNoTI, but if it is an effect of TI, we expect it to show positive effects of the contrast (stimTI – shamTI) – (stimNoTI – shamNoTI). Our sample size of 8 subjects is too small to draw strong conclusions from, but we see a significant positive effect of (stimTI – shamTI) – (stimNoTI – shamNoTI) in parts of temporal gyrus (figure S1 in the supplementary material). This is consistent with the notion that the activation in the parahippocampal gyrus might be an effect of TI itself and not of high-frequency currents or off-target tonic activation.

Further, is it necessary to optimize stimulation set-up for individual subjects? Our results show that the variability in peaks of predicted TI across the subjects is low compared to variability in peaks of BOLD. This suggests that individualized optimization of the electrode locations may yield only minimal improvement in the spatial precision of the resulting BOLD signal. Still, these observations could result from invalidity of the approximations of FEM for TI prediction (e.g. assuming isotropy) or from differences across the subjects in the relationship between TI and BOLD activation. Thus, it seems beneficial to perform a systematic inquiry of how much the differences in tissue conductance affect the predicted TI as a part of future research direction. It would also be beneficial to develop models based on structural connectivity data to make better predictions of the BOLD activation given the predicted TI. These future research efforts would be critical in satisfactorily resolving this question of the utility of individualized optimization.

Overall, the results suggest that TI may be a safe and effective way to modulate BOLD signals in the human brain, especially in the deeper regions typically out of reach of other methods.

## Acknowledgments

Supported by an Indiana University Grand Challenge in Addiction grant.

