## Supplementary material for "Temporal interference electrical neurostimulation yields fMRI BOLD activation in humans"

**Section A**: Preliminary results of (StimTI – ShamTI) – (StimNoTI – ShamNoTI) contrast

Preliminary effects of temporal interference (TI) as distinct from effects of electric fields of high frequency carrier current. In the “NoTI” condition, the frequencies of both electrode pairs are the same, i.e. 2000Hz, thus eliminating the temporal interference pattern. These results are preliminary given that there were only 8 subjects in this condition.

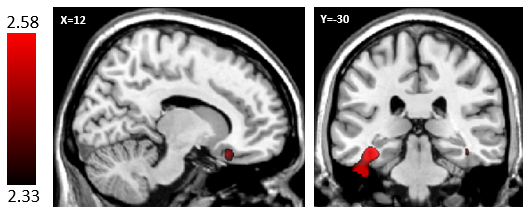

Figure S1: Positive effects of (StimTI – ShamTI) – (StimNoTI – ShamNoTI) contrast

Table S1: Positive effects of (StimTI – ShamTI) – (StimNoTI – ShamNoTI) contrast. Cluster-defining threshold = 0.001; Cluster extent threshold = 5

|  |  |  | Peak MNI coordinates | | |  |  |
| --- | --- | --- | --- | --- | --- | --- | --- |
| Region | **Laterality** | **Cluster Size** | **X** | **Y** | **Z** | **Max stat t** | **P Cluster Corrected** |
| Middle temporal gyrus, inferior temporal gyrus, fusiform gyrus | Left | 775 | -52 | 0 | -24 | 18.62 | 0.04 |

The cluster in right orbito-frontal gyrus has peak at [18,28,-30] (z = 3.07, n = 8 subjects) but is not significant upon whole brain cluster correction or small-volume correction using the mask in figure 2B.

There are no significant negative effects of (StimTI – ShamTI) – (StimNoTI – ShamNoTI) contrast.

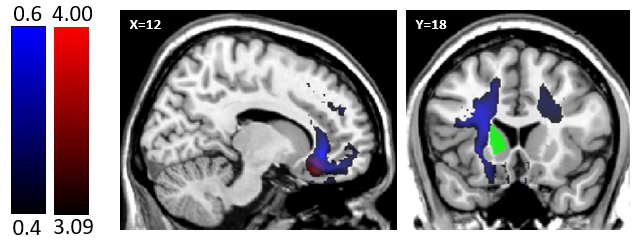

Figure S2: TI in target region was not the highest. Blue: Voxels with TI^*^>=0.4. TI^*^ refers to FEM simulation values in all-subject’s TI image (as described in section 2.2 and S – F.7); Red: Cluster in frontal cortex showing significant fMRI BOLD activation for contrast stimTI – ShamTI after small volume correction (See figure 2B for this mask); Green: Our initial target region.

**Section B**: Predicted TI values across the subjects are highly correlated.

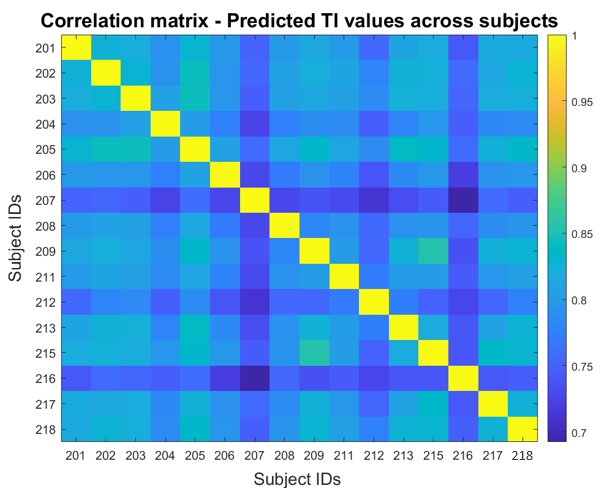

Figure S3: Correlation matrix of predicted TI values across subjects. A particular tile shows the correlation between the predicted TI values for the voxels in MNI space of subjects corresponding to the row and the column.

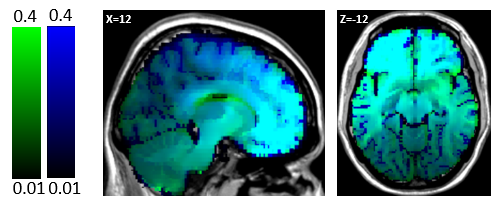

Figure S4: TI prediction for the reference subject (Green) overlaid on mean of the TI predictions for the study participants (Blue).

The predicted TI values for the reference subject and the average of the predicted TI values across the subjects are overlayed on each other in figure S2. Supplementary figure S3 shows the correlation matrix for the predicted TI values across the subjects, with r values > 0.70 (p < 0.001) across all subject pairings.

**Section C**: What is the relationship between the predicted TI field and the BOLD signal? (Main article – section 3.5 continued)

The specific functional relationship between TI* and BOLD signals seemed to involve a positive correlation, but the details remain to be clarified. We asked the following question: at the voxel level, which function of TI best predicts the observed BOLD activation? This analysis was performed only for the voxels in gray matter.

For every subject, we considered that the BOLD signal may be a nonlinear function of the TI field, f(TI). We considered various forms of f(TI) such as polynomial, exponential, logistic function, etc and found the optimal parameters that minimize the sum of squared errors. We compared a total of 13 models in this manner using the Akaike Information Criteria (AIC). We fit each model individually to the subjects and found that for all subjects the logistic model of TI performs the best in predicting BOLD contrast values. The results of this model comparison are provided in tables S2 and S3.

The parameters of this model were not significant in one-sample t-test across the subjects, which prevents us from identifying a single set of parameter values that can be used to predict BOLD activation. We noticed that there were some subjects that differed qualitatively in their relation between TI and BOLD contrast values.

We performed hierarchical clustering of the subjects using the hamming distance between the best fitted parameters of logistic model and identified two sub-groups which show qualitatively different relation between predicted TI and BOLD contrast values based on this model (Figure S5A). For this, we first identified the parameters that determine the direction of variation of BOLD with TI values. We refer to these as parameters of interest. Consider the logistic model in equation C.1.

$y \sim c1 + \frac{c2}{(1+ e^{-(c3*x+c4)})}$	Eq. (C.1)

The signs of the parameters c2 and c3 determine the direction of change in y with some change in x. Various possible combinations of the signs of the parameters of interest were uniquely coded and this code assigned to the subjects. The hamming distance between these were used to cluster the subjects.

The larger subgroup (11 subjects) corresponds to parameter values such that the model predicts higher BOLD contrast value when TI is higher while the smaller sub-group (5 subjects) corresponds to the parameter values that the model predicts lower BOLD contrast values when the TI is higher.

We similarly also performed hierarchical clustering of the subjects using the best fitted parameter values of top five models from the model comparison, taken together. For this, each subject was represented as a row-vector with each column representing the qualitative relationship between the TI values and the model-predicted BOLD activation for a particular model. Subjects were clustered using the hamming distance between these row-vectors. Please refer to figure S5B to see the results of this which also led to the same two sub-groups as identified with the clustering performed with only the parameters of logistic model.

The clustering results raised concerns about whether our main finding in figure 2 is driven by the sub-group that showed a positive relation between predicted TI and BOLD. We performed the one-sample t-test of our fMRI results separately for these sub-groups (figures S6 and S7). The effect for bigger group has MNI peak at {-30, 42, 6} (t=5.56, p = 0.008, cluster size = 979). The effect for the smaller group has the MNI peak at {24, 30, -30} (t = 12.52, p-value = 0.345, cluster size = 36). Both these results are obtained after small volume correction using the mask shown in figure 2C. The frontal clusters in these sub-groups appear to be localized in different lobes. Although the effect for the bigger group is stronger, we found that both sub-groups show effect of stimTI – shamTI contrast in the frontal lobe even when analyzed separately. The weaker effect observed in smaller groups could be owing to its smaller size. It must be noted that the predicted TI values are high (TI*>0.6) in both lobes as can be seen in figure 2C.

Possible reasons for why the subjects are qualitatively different in their direction of variation with TI may be individual differences in functional connectivity leading to differential TI-induced BOLD activation. It is also possible that assumptions of head-models such as isotropy may not hold well for some subjects leading to inaccurate predictions of TI. It may also be possible that these subjects in one group had a different cognitive experience of the TIS than the subjects in other groups. We had asked subjects about any distresses or discomfort throughout the stimulation session in the form of the stimulation experience scale (SES) as reported in the main article. These scales did not reveal anything significantly different between these groups, but it is possible that the differences lie in an experience that our questionnaires did not capture.

Despite differences in relation between BOLD contrast values and predicted-TI values across the brain, both these sub-groups show the positive effect of stimTI – shamTI contrast in the frontal lobe where the TI was predicted to be high (TI* > 0.6).

Table S2: Results of model comparison of different models of TI whose parameters were optimized for predicting the BOLD contrast values stimTI – shamTI at the level of voxels. Model comparison is based on aggregate AIC.

| Model Number | Model | Aggregate AIC |
| --- | --- | --- |
| 1 | $y \sim c1 + c2*x$ | $-1.6269* {10}^{6}$ |
| 2 | $y \sim c1 + c2*x^{2}$ | $-1.5974* {10}^{6}$ |
| 3 | $y \sim c1 + c2*x^{3}$ | $-1.5584* {10}^{6}$ |
| 4 | $y \sim c1 + c2*x+ c3*x^{2}$ | $-1.6532* {10}^{6}$ |
| 5 | $y \sim c1+ c2*x^{2}+ c3*x^{3}$ | $-1.6359* {10}^{6}$ |
| 6 | $y \sim c1 + c2*x+ c3*x^{2}+ c4*x^{3}$ | $-1.6663* {10}^{6}$ |
| 7 | $y \sim c1 + c2*\sqrt{x}$ | $-1.6356* {10}^{6}$ |
| 8 | $y \sim c1 + c2*\frac{1}{x}$ | $-1.6334* {10}^{6}$ |
| 9 | $y \sim c1 + c2*\frac{1}{x^{2}}$ | $-1.6010* {10}^{6}$ |
| **10** | $\boldsymbol{y \sim c}\boldsymbol{1 +}\frac{\boldsymbol{c}\boldsymbol{2}}{\boldsymbol{(1+}\boldsymbol{e}^{\boldsymbol{-(c}\boldsymbol{3*x+c}\boldsymbol{4)}}\boldsymbol{)}}$ | $\boldsymbol{-1.7467*}\boldsymbol{10}^{\boldsymbol{6}}$ |
| 11 | $y \sim c1 + {c2*e}^{c3*x}$ | $-1.6321* {10}^{6}$ |
| 12 | $y \sim c1 + c2*log(x+c3)$ | $-1.6412* {10}^{6}$ |
| 13 | $y \sim c1 + \frac{c2*x}{(c3+x)}$ | $-1.6456* {10}^{6}$ |

In table S2,

y = Value of fMRI BOLD contrast stimTI – shamTI at the level of voxels

x = Value of the predicted TI at the level of voxels. These predictions were made by constructing a finite element model of the subjects using their structural as explained in the methods section in the main article.

Here, c_i_ refers to model parameters where ‘i’ is a subscript going from 1 to number of parameters in the models. Parameters of a particular model were found such that they minimized the sum of squared errors (SSE) in predicting the BOLD contrast values. Each model was individually fit to a subject and its SSE noted.

Aggregate AIC: For a model, its SSE for each subject’s data was summed to get an aggregate SSE which was used to find the AIC of the model. In finding AIC, the number of parameters were taken as number of parameters in the model times the number of subjects because the model parameters were individually fit to the subjects.

Arranging the models from best to worst based on their aggregate AIC:

10	6	4	13	12	5	7	8	11	1	9	2	3

Table S3:Results of model comparison of different models of TI for predicting the stimTI-shamTI BOLD contrast values at the level of voxels. Model comparison is performed for each subject based on their AIC. Models are referred by their number as given in table S2.

| Subject | Best |  |  |  |  |  |  |  |  |  |  |  | Worst |
| --- | --- | --- | --- | --- | --- | --- | --- | --- | --- | --- | --- | --- | --- |
| 1 | 10 | 6 | 11 | 13 | 5 | 8 | 4 | 9 | 2 | 1 | 12 | 3 | 7 |
| 2 | 10 | 6 | 4 | 9 | 8 | 13 | 12 | 5 | 7 | 1 | 11 | 2 | 3 |
| 3 | 10 | 6 | 4 | 5 | 13 | 12 | 11 | 7 | 1 | 8 | 2 | 9 | 3 |
| 4 | 10 | 6 | 11 | 13 | 8 | 4 | 12 | 7 | 1 | 5 | 9 | 2 | 3 |
| 5 | 10 | 6 | 5 | 1 | 4 | 11 | 12 | 13 | 2 | 7 | 8 | 3 | 9 |
| 6 | 10 | 4 | 6 | 5 | 11 | 7 | 13 | 12 | 1 | 8 | 9 | 3 | 2 |
| 7 | 10 | 6 | 11 | 13 | 4 | 12 | 7 | 8 | 1 | 5 | 2 | 9 | 3 |
| 8 | 10 | 6 | 11 | 8 | 13 | 4 | 12 | 9 | 7 | 5 | 1 | 2 | 3 |
| 9 | 10 | 6 | 11 | 13 | 12 | 4 | 7 | 1 | 5 | 8 | 2 | 3 | 9 |
| 10 | 10 | 6 | 11 | 4 | 13 | 12 | 7 | 5 | 1 | 2 | 3 | 8 | 9 |
| 11 | 10 | 6 | 4 | 8 | 9 | 13 | 11 | 12 | 5 | 7 | 1 | 2 | 3 |
| 12 | 10 | 6 | 4 | 8 | 11 | 13 | 9 | 5 | 12 | 7 | 1 | 2 | 3 |
| 13 | 10 | 5 | 6 | 4 | 11 | 13 | 12 | 1 | 7 | 2 | 3 | 8 | 9 |
| 14 | 10 | 6 | 11 | 4 | 8 | 13 | 12 | 9 | 5 | 7 | 1 | 3 | 2 |
| 15 | 10 | 6 | 4 | 11 | 13 | 12 | 7 | 5 | 1 | 8 | 2 | 3 | 9 |
| 16 | 10 | 6 | 5 | 2 | 4 | 1 | 13 | 11 | 12 | 7 | 8 | 3 | 9 |

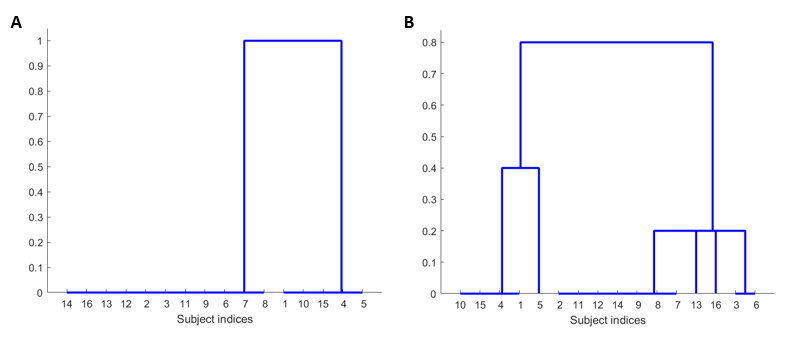

Figure S5: Clustering of subjects based on the qualitative relation between predicted-TI and stimTI – shamTI BOLD contrast values as indicated by the best-fit parameter values of: A – Best (logistic) model and B -top five models, from the model comparison.

Contrast stimTI – ShamTI – for the two subgroups

- Bigger sub-group (Subjects 2, 3, 6, 7, 8, 9, 11, 12, 13, 14, and 16)

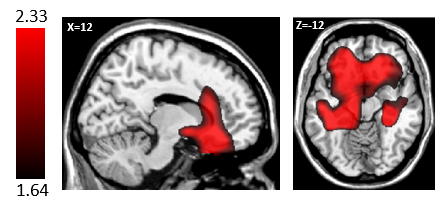

Figure S6: Positive effects of (StimTI – ShamTI) contrast in bigger sub-group

- Smaller sub-group (Subjects 1, 4, 5, 10, and 15)

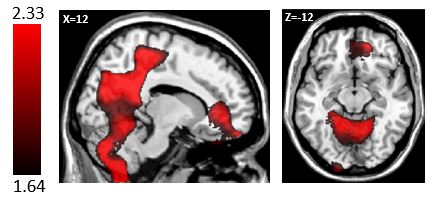

Figure S7: Positive effects of (StimTI – ShamTI) contrast in smaller sub-group

**Section D**: Peak MNI coordinates of BOLD activation; Overlap of all-subjects TI image and average electrode locations

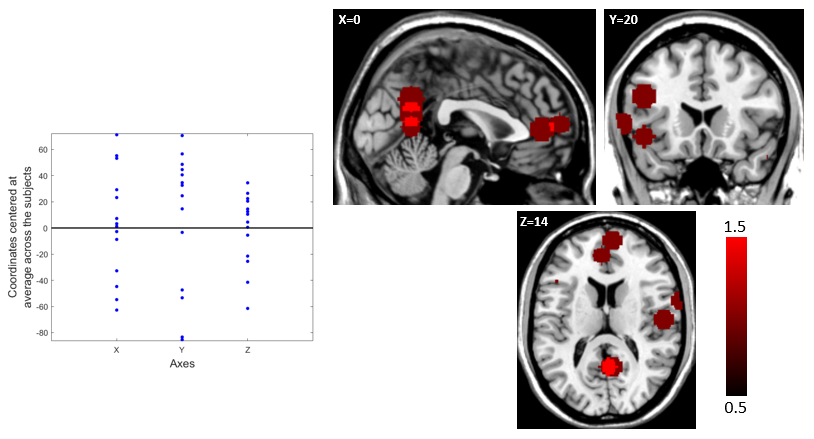

Figure S8: Variability in peak MNI coordinates for BOLD contrast values for (stimTI – shamTI) across subjects. Left: Plot shows the peak MNI coordinates of the BOLD contrast for subjects. The X, Y, Z coordinates are separately centered at their means across the subjects and are shown as blue dots. Right: Red shows the overlap of 10 mm spheres around the peak MNI coordinates of the subjects.

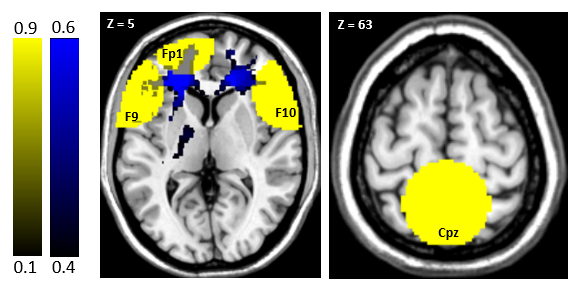

Figure S9: All-subjects TI image (blue) overlapped with the masks around the average (across subjects) positions of the electrodes. Predicted TI is higher in vicinity of the electrodes F9, F10, and Fp1 (Left) than in that of electrode Cpz (Right). The maximum of the values of all-subjects TI image within the mask is lowest in case of Cpz. Mean of the values of all-subjects TI image within the mask is lowest for both Cpz and F9 with the one for F9 being marginally lower than Cpz.

**Section E**: Stimulation Experience Scale (SES)

Table S4: Raw ratings of all subjects on SES after every block. SES is stimulation experience scale that they were administered to collect their self-report on the various aspects of their experience of the block on a scale of 1-5 (lowest to highest).

| Subject | Block | Stress | Anxiety | General anxiety | Ringing in ears | Headache | | Itchiness | | Nausea | | Tingling on the skin | | Numbness on the skin | | Muscle pain | | Muscle contractions | Total neurostimulation experience score^*^ |
| --- | --- | --- | --- | --- | --- | --- | --- | --- | --- | --- | --- | --- | --- | --- | --- | --- | --- | --- | --- |
| 1 | Sham TI | 1 | 1 | 1 | 1 | 1 | 1 | | 1 | | 1 | | 1 | | 1 | | 1 | | 11 |
|  | Stim TI | 1 | 1 | 1 | 1 | 1 | 1 | | 1 | | 1 | | 1 | | 1 | | 1 | | 11 |
|  | Sham TI | 1 | 1 | 1 | 1 | 1 | 1 | | 1 | | 1 | | 1 | | 1 | | 1 | | 11 |
|  | Stim TI | 1 | 1 | 1 | 1 | 1 | 1 | | 1 | | 1 | | 1 | | 1 | | 1 | | 11 |
| 2 | Sham TI | 1 | 1 | 1 | 1 | 1 | 1 | | 1 | | 1 | | 1 | | 1 | | 1 | | 11 |
|  | Stim TI | 1 | 1 | 1 | 1 | 1 | 1 | | 1 | | 1 | | 1 | | 1 | | 1 | | 11 |
| 3 | Stim TI | 2 | 2 | 1 | 1 | 1 | 1 | | 1 | | 1 | | 1 | | 1 | | 1 | | 13 |
|  | Sham TI | 1 | 1 | 1 | 1 | 1 | 1 | | 1 | | 1 | | 1 | | 1 | | 1 | | 11 |
|  | Stim TI | 1 | 1 | 1 | 1 | 1 | 1 | | 1 | | 1 | | 1 | | 1 | | 1 | | 11 |
|  | Sham TI | 1 | 1 | 1 | 1 | 1 | 1 | | 1 | | 1 | | 1 | | 1 | | 1 | | 11 |
| 4 | Stim TI | 2 | 2 | 1 | 1 | 1 | 1 | | 1 | | 1 | | 1 | | 1 | | 1 | | 13 |
|  | Sham TI | 1 | 1 | 1 | 1 | 2 | 1 | | 1 | | 1 | | 1 | | 1 | | 1 | | 12 |
|  | Stim TI | 1 | 1 | 1 | 1 | 2 | 1 | | 1 | | 1 | | 1 | | 1 | | 1 | | 12 |
|  | Sham TI | 1 | 1 | 1 | 1 | 2 | 1 | | 1 | | 1 | | 1 | | 1 | | 1 | | 12 |
| 5 | Sham TI | 1 | 1 | 1 | 1 | 1 | 1 | | 1 | | 1 | | 1 | | 4 | | 1 | | 14 |
|  | Stim TI | 1 | 1 | 1 | 1 | 1 | 1 | | 1 | | 1 | | 1 | | 1 | | 1 | | 11 |
|  | Sham TI | 1 | 1 | 1 | 1 | 1 | 1 | | 1 | | 1 | | 1 | | 4 | | 1 | | 14 |
|  | Stim TI | 1 | 1 | 1 | 1 | 1 | 1 | | 1 | | 1 | | 1 | | 4 | | 1 | | 14 |
| 6 | Sham TI | 2 | 1 | 1 | 1 | 1 | 1 | | 1 | | 1 | | 1 | | 1 | | 1 | | 12 |
|  | Stim TI | 1 | 1 | 1 | 1 | 1 | 1 | | 1 | | 1 | | 1 | | 1 | | 1 | | 11 |
|  | Sham TI | 1 | 1 | 1 | 1 | 1 | 1 | | 1 | | 1 | | 1 | | 1 | | 1 | | 11 |
|  | Stim TI | 1 | 1 | 1 | 1 | 1 | 1 | | 1 | | 1 | | 1 | | 1 | | 1 | | 11 |
| 7 | Stim TI | 1 | 2 | 1 | 1 | 1 | 1 | | 1 | | 1 | | 1 | | 1 | | 1 | | 12 |
|  | Sham TI | 1 | 1 | 2 | 1 | 1 | 1 | | 1 | | 2 | | 3 | | 1 | | 1 | | 15 |
|  | Stim TI | 1 | 2 | 2 | 1 | 1 | 1 | | 1 | | 1 | | 2 | | 1 | | 1 | | 14 |
|  | Sham TI | 1 | 2 | 1 | 1 | 1 | 1 | | 1 | | 2 | | 2 | | 1 | | 1 | | 14 |
| 8 | Stim TI | 2 | 2 | 1 | 1 | 1 | 1 | | 1 | | 1 | | 1 | | 1 | | 1 | | 13 |
|  | Sham TI | 2 | 2 | 2 | 1 | 1 | 1 | | 1 | | 1 | | 1 | | 1 | | 1 | | 14 |
|  | Stim TI | 2 | 2 | 1 | 1 | 1 | 1 | | 1 | | 1 | | 1 | | 1 | | 1 | | 13 |
|  | Sham TI | 2 | 2 | 1 | 1 | 1 | 1 | | 1 | | 1 | | 1 | | 1 | | 1 | | 13 |
| 9 | Sham TI | 1 | 1 | 1 | 1 | 1 | 1 | | 1 | | 1 | | 1 | | 1 | | 1 | | 11 |
|  | Stim TI | 1 | 1 | 1 | 1 | 1 | 1 | | 1 | | 1 | | 1 | | 1 | | 1 | | 11 |
| 10 | Sham TI | 1 | 1 | 1 | 1 | 1 | 1 | | 1 | | 1 | | 1 | | 1 | | 1 | | 11 |
|  | Stim TI | 1 | 1 | 1 | 2 | 2 | 1 | | 1 | | 2 | | 1 | | 1 | | 1 | | 14 |
| 11 | Sham TI | 1 | 1 | 1 | 1 | 1 | 1 | | 1 | | 1 | | 1 | | 1 | | 1 | | 11 |
|  | Stim TI | 1 | 1 | 1 | 1 | 1 | 1 | | 1 | | 1 | | 1 | | 1 | | 1 | | 11 |
| 12 | Stim TI | 2 | 2 | 1 | 1 | 1 | 1 | | 1 | | 1 | | 1 | | 1 | | 1 | | 13 |
|  | Sham TI | 1 | 1 | 1 | 1 | 1 | 1 | | 1 | | 1 | | 1 | | 1 | | 1 | | 11 |
| 13 | Stim TI | 2 | 2 | 1 | 1 | 1 | 1 | | 1 | | 1 | | 1 | | 1 | | 1 | | 13 |
|  | Sham TI | 2 | 1 | 1 | 1 | 1 | 1 | | 1 | | 1 | | 1 | | 1 | | 1 | | 12 |
| 14 | Stim TI | 1 | 1 | 1 | 1 | 1 | 1 | | 1 | | 1 | | 1 | | 1 | | 1 | | 11 |
|  | Sham TI | 1 | 1 | 3 | 1 | 1 | 1 | | 1 | | 1 | | 1 | | 1 | | 1 | | 13 |
| 15 | Stim TI | 1 | 1 | 1 | 1 | 1 | 1 | | 1 | | 1 | | 1 | | 1 | | 1 | | 11 |
|  | Sham TI | 1 | 1 | 2 | 1 | 1 | 2 | | 1 | | 2 | | 1 | | 1 | | 4 | | 17 |
| 16 | Stim TI | 4 | 2 | 1 | 1 | 1 | 1 | | 1 | | 1 | | 1 | | 1 | | 1 | | 15 |
|  | Sham TI | 2 | 2 | 2 | 3 | 1 | 1 | | 1 | | 1 | | 1 | | 1 | | 1 | | 16 |

^*^Lowest possible value is 11 and highest is 55 with higher score indicating more discomfort.

Table S5: Number of blocks with the rating of 2 or more in each of the categories in the SES scale. Rating ‘1’ – No discomfort in the given category; Rating ‘5’ – High discomfort in the given category. Each of stim-TI and sham-TI conditions had 23 blocks in total across the subjects.

| Block | Stress | Anxiety | General anxiety | Ringing in ears | Headache | Itchiness | Nausea | Tingling on the skin | Numbness on the skin | Muscle pain | Muscle contractions |
| --- | --- | --- | --- | --- | --- | --- | --- | --- | --- | --- | --- |
| Stim TI | 7 | 3 | 1 | 1 | 2 | 0 | 0 | 1 | 1 | 1 | 0 |
| Sham TI | 5 | 8 | 4 | 0 | 2 | 1 | 0 | 3 | 2 | 2 | 1 |

**Section F**: More details about the study

**F.1 Participants and recruitment criteria** (Main article – Section 2.1 continued)

A total of 16 subjects (Age: mean = 28.9 years, standard deviation = 10; Sex: 10 females and 6 males) were recruited via Craigslist, the Indiana University Classifieds, and flyers placed throughout Bloomington, Indiana. Subjects were required to be between 18 and 50 years of age, weigh less than 440 pounds, have at least a 6^th^ grade education, and be able to speak and read English. Subjects were ineligible if they were taking ADHD medication, or if they had migraine headaches, epilepsy, a seizure disorder, metal implants in their head or scalp, any neurological disease, symptoms of psychosis, or any other conditions that would make it unsafe for them to enter the MRI scanner.

All subjects participated in TI conditions while only the last 8 participated in no-TI condition (See section 2.5 for description of ‘no-TI’).

**F.2** **Neurostimulation device and apparatus** (Main article - Section 2 continued)

The Soterix Temporal Interference electrical neurostimulation device (Soterix Medical, Inc.) was used to deliver TI stimulation. The TI device contains a mechanism that constantly monitors the amount of current administered via the electrodes and is rated to deliver up to 2mA per electrode pair. The device has the following characteristics. It is powered by four rechargeable D-cell batteries that provide the energy to carry out neurostimulation. There are two electrode pairs, with a total of 4 electrodes, that are connected to neurostimulation device on one end and to the subject’s scalp on the other end. At 2 mA current, the stimulation results in a maximum current density of 0.002 A/cm2. At 50 KΩ scalp impedance the total power dissipated by an electrode pair is I^2^R = 200 mW, which is comparable to the power emitted by a cell phone. Each electrode pair generates a sinewave alternating current, with no DC component.

**F.3 Study procedure** (Main article – Section 2.1 continued)

All procedures involving human subjects were approved by the IRB of Indiana University. After providing informed consent, subjects were asked to complete measures of mood, working memory, and physical sensations, including a newly developed Stimulation Experience Scale (SES). The SES measured the levels of discomfort as well as presence of any potential unusual sensory experiences/suicidal ideation. Fasteners for all 4 MRI-compatible carbon electrodes were placed into designated locations on an electrode EEG cap, at F9 and F10 for one of the electrode pairs, and at FP1 and CPz for the other pair. Electromedical gel was used to secure electrodes into the holders such that the gel made the connection between the electrodes and the scalp.

The Impedance of the connection was checked and brought to less than or equal to 50 kΩ by adding more gel if required before beginning the scanning. Inside the MRI scanner, subjects were administered neurostimulation as explained in section 2.5.

The SES was administered several times throughout the course of the session, including before, during, and after the stimulation procedure. The ratings provided by the subjects for the SES are provided in table S4 in supplementary materials. Subjects also completed repeated measures of their mood, memory, and physical sensations both immediately after as well as 10 and 30 minutes after stimulation. Subjects received a follow-up call between 2 and 15 days after the study in order to query the emergence of any cognitive changes or other potential adverse effects.

**F.4** **Detailed experiment design** (Main article – section 2.3 continued)

We conducted four 8-min blocks with alternating blocks of active and sham stimulation such that there were two blocks of active stimulation and two blocks of sham stimulation. The active stimulation block consisted of 120 seconds of stimulation (which we refer to as ‘ON’) alternating with 120 seconds of no stimulation where zero current was passed through the electrodes (which we refer to as ‘OFF’). The ON period consisted of 30 seconds of the current ramping up linearly from 0 mA to 2 mA across the two electrodes in both pairs of electrodes and 90 seconds when the current across both pairs remained at 2 mA. The choice of 2 mA as the maximum current amplitude was used as previous studies using TI in humans have also used same current magnitude and suggest this to be a safe choice (Ma et al., 2022; Zhu et al., 2022). The OFF period consisted of 30 seconds of ramping down from 2 mA to 0 mA and 90 seconds of 0 mA through both pairs of electrodes. The time taken for ramp-up and ramp-down was a characteristic of the neurostimulation controller and served to minimize unpleasant scalp sensations that would otherwise result from rapid current changes. The choice of 120 seconds as the duration for a single uninterrupted stimulation (‘ON’) and no-stimulation (‘OFF’) was driven by several considerations. First, to avoid the effect of ON vs OFF being lost in linear detrending of the fMRI data in preprocessing, it was necessary to at least have two Ons and two OFFs per block. Second, the total duration of one block was 8 minutes, and time required for ramping up and down the current was 60 seconds. The duration of the ON period required at least 60 seconds for the current to ramp-up and down as well as sufficient time for the current to stay at its maximum value so that the effect of active – sham could be observed. This weighed against having more than two ON periods. Thus, we ran two ON periods of two minutes each, alternating with two OFF periods. The order of whether ON vs. OFF periods started first was counterbalanced across runs and subjects.

The sham block had alterations between ON and OFF just like the active block, however, the ON in the sham did not have the 60 seconds of stimulation between ramp-up and ramp-down. It only consisted of 30 seconds of ramping up from 0 to 2 mA followed by immediate ramp down from 2 to 0 mA. The sham block controlled for the sensations on the scalp that the subjects feel/may feel when being stimulated, which may occur during the ramp up.

A particular block could either begin with an ‘ON’ or an ‘OFF’, and we refer to blocks as ON-block or an OFF-block depending on what they began with. The active ON-block and its corresponding sham ON-block (figure 1 A,B) were conducted consecutively, in any order. Same was done in case of active OFF-block, and its corresponding sham OFF-block. Further, no two sham blocks or two active blocks, irrespective of whether they were ON-or OFF-blocks, could be together.

Further a block could either be a TI-block or a no-TI-block. In the TI condition, the currents through two pairs of electrodes were administered at frequencies, 2000 Hz and 2020 Hz so that the interference pattern of their induced electric fields was expected to have a low frequency component of 20 Hz. In the no-TI-condition, both the currents were administered at 2000 Hz, so that their interference pattern would have no low frequency component, that is, there was ‘no TI’. The no-TI-condition controlled for any effects of high frequency currents on the fMRI BOLD activation (Mirzakhalili et al., 2020) thus allowing us to identify BOLD effects of TI as distinct from BOLD effects that may be due to the individual high frequency stimulation.

Subjects were run on two different variations of the above:

Design 1: For a subject, a session had both ON- and OFF-blocks. Further, the session could either begin with stimulation or a sham block. All the blocks were TI blocks. These were counterbalanced across subjects such that for a subject, a session could be any of the four possibilities. One example of a session for a subject is: ON-stim-TI 🡪 ON-sham-TI 🡪 OFF-stim-TI 🡪 OFF-sham-TI.

Design 2: We also collected a smaller set of data for a slightly different design. In this, for a subject, a session had both TI and no-TI conditions. Further, the session could either begin with stimulation or a sham block. All the blocks for a given subject were either ON or OFF. These were counterbalanced across subjects such that for a subject, a session could be any of the eight possibilities. One example of a session for a subject in this design is: ON-stim-TI 🡪 ON-sham-TI 🡪 ON-stim-no-TI 🡪 ON-sham-no-TI.

**F.5** **fMRI acquisition and preprocessing** (Main article - section 2 continued)

The fMRI acquisition and preprocessing methods have been described previously (Modak et al., 2021). A Siemens 3 Tesla TIM Trio MRI scanner at the imaging research facility at the Psychological and Brain Sciences department at Indiana University Bloomington was used to collect functional MRI data using Echo-planar imaging (EPI) pulse sequences. The functional and structural scans were collected using a 32-channel head coil, and with axial slices at an angle of 30^o^ with the line connecting anterior commissure and posterior commissure. The slice angle was chosen to improve the signal to noise ratio in the orbitofrontal cortex, where we expected to find signal (Deichmann et al., 2003). The T2* weighted functional scans were collected with a TR = 2000 ms, TE = 25 ms, flip angle = 70^o^, and a 64 x 64 acquisition matrix. For each run, 240 volumes were collected with each volume consisting of 35 slices of the thickness of 3.8 mm.

T1 weighted functional scans were collected with a TR = 1800 ms, TE = 2.7 ms, flip angle = 9^o^, and a 256 x 256 acquisition matrix. In this case, 160 slices were collected with a thickness of 1 mm.

SPM12 was used to preprocess the data. Spike correction was performed using 3dDespike in AFNI, and the slice timing correction was performed in SPM12. For every subject, the functional scans were first realigned, then co-registered to the structural scan, and then normalized to the standard Montreal Neurological Institute (MNI) space. Spatial smoothing was performed once using an 8 mm^3^ kernel and later a 20 mm^3^ full-width-at-half-maximum (FWHM) kernel on the premise that the larger smoothing kernel would allow us to see results with greater spatial variability at the expense of statistical power in any one region.

We report the results from the GLM analysis done for the data with 20 mm smoothing. With a smaller smoothing kernel (8mm), the effect of stimulation (StimTI – ShamTI) was seen as distributed clusters in the target area across the subjects leading to no one region showing significant effect across the subjects.

**F.6 Finding optimal electrode locations** (Main article – Section 2.2 continued)

The optimal locations for the four electrodes (that is 2 pairs of electrodes) were identified in the EEG 10-10 electrode system with the help of the SimNIBS software (Saturnino, Puonti, et al., 2019) which was used to create the finite element model (FEM) of the brain and find the induced electric field due to the current through the electrodes. In finding the induced electric field, we assumed isotropy in conductance. Structural images from one subject which we refer to as the **reference subject** from here on were used to first create a head model using the headreco utility in SimNIBS. The optimal electrode positions were found for the reference subject in a manner specified below, and these same electrode locations were used to stimulate all subjects in this study.

The optimization was performed using SimNIBS to maximize the focality of the induced electric field at the target location (that is, to maximize the electric fied in the region within 10 mm of the coordinates of the target location identified using the neuromorphometrics atlas, Neuromorphometrics, Inc., Winthrop, MA, USA). This optimization was with the constraints that the total current administered across all four electrodes could at maximum be 4 mA; the total number of active electrodes could at maximum be four; the current through any one electrode could at maximum be 2 mA, and the intensity of the electric field in the target region is 0.4 V/m. This corresponds to the optimization problem 4 in Saturnino et al. (2019) as implemented in SimNIBS. The electrodes that were part of the solution of the above problem were used to construct multiple two-pairs of electrodes, that is, multiple combinations consisting of four electrodes such that in each combination, there are two pairs. To find the optimal electrode locations, we consider all possible combinations of electrode locations. For a particular combination, the induced electric field due to 2 mA passed through each of the two pairs of electrodes was found. Further, temporal interference resulting from current passed through these two pairs was found. Let E1 represent the amplitude of the magnitude of electric field at every element in the mesh (head model) due to 2 mA current through one pair of electrodes and E2 represent the same for the other pair of electrodes. The maximum amplitude modulation of the interference pattern is given by the following equation (Grossman et al. (2017)):

${TI}_{amp}=\left\{ \begin{aligned} 2E_{2}, if E_{2}<E_{1}cos(\alpha) \\ \frac{(2\left| \vec{E_{2}}\times(\vec{E_{1}}- \vec{E_{2}}) \right|)}{\left| \vec{E_{1}}- \vec{E_{2}} \right|}, &otherwise \end{aligned} \right.$	Eq (F.1)

Here, E_2_<E_1_ and α < 90 degrees, and note that the TI field amplitude is limited to the weaker of the two fields E_1_ and E_2_.

This amplitude was taken as the measure of TI such that wherever TI_amp_ was high TI was considered to be high. These values were normalized to the MNI space.

This was repeated for each of the possible combinations of electrode locations. The next task was to choose one combination from these combinations that best served our purpose. This was done using the following objective function, which aims to maximize TI to the target region and minimize it elsewhere:

$$Maximize TI_{fm} where TI_{fm}=6*M1+ 5*M2+3*M3+4*M4+5*M5$$

Here,

TI_fm_ is a measure of focality that we maximized.

M1 is the average (across the non-target regions) of the ratio of difference between the mean of the TI in the target and non-target regions expressed as the percentage of mean of the TI in the non-target regions.

M2 is same as M1, except that it does not correspond to mean TI in the regions but to the 99-percentile value of TI in the regions.

M3 is a measure of entropy such that higher values indicate lower entropy in target region relative to the non-target region, i.e. higher uniformity in magnitude of the stimulation amplitude within the target region.

M4 represents the average ratio (across non-target regions) of the difference between X1 and X2 as a percentage of X2 where X1 is average of the ratio of TI values in the target region and 99 percentile TI value of the whole brain and X2 is same as X1 but for non-target regions.

M5 is the number of voxels in the target region that have a TI > 0.2 V/m expressed as the percentage of the average number of the voxels in the non-target regions that have a TI > 0.2 V/m.

This weighting scheme was chosen as a reasonable approach to optimizing the targeting, although other approaches are possible and may yield similar results.

To calculate the TI_fm_, we used the neuromorphometrics atlas as mentioned above, but before parcellating, we first masked out the white matter using the gray matter mask of the reference subject generated during segmentation. This way, only the TI in the voxels in gray matter contributed to the TI_fm_ of different regions.

The combination, that is, the two pairs of electrodes which had the maximum TI_fm_ for the reference subject, were used in the stimulation protocol for all subjects.

**F.7 Individualized finite element analysis (FEA) and all-subjects TI image** (Main article – Section 2.2 continued)

Along with the fMRI data, we also collected the structural image of the subjects. For every subject, we used this structural image to construct a head model and perform the FEA to find the expected temporal interference field magnitude at every voxel in the subject’s brain. This was necessary to accurately predict the TI across subjects, as our optimization of the stimulation setup was done only for the reference subject, but every subject’s head is different in terms of their head size and brain segmentation leading to differences in predicted TI from that of the reference subject. The locations of the electrodes in subject space were visually inspected from the structural and were used in finding the induced electric fields and subsequently the magnitude of temporal interference (TI). This was done to minimize any inaccuracies in TI prediction due to the electrodes getting differentially shifted by the stretching of the EEG cap. FEA yielded the electric fields induced by both pairs of electrodes which were then used to estimate the TI_amp_ in a manner described in section 2.4.1. These estimates were normalized to 2mm MNI space.

To predict the temporal interference effect on BOLD signals that we would expect to see across the subjects, we created an image of the mean minus two standard deviations of the predicted TI values across the subjects for every voxel in MNI space. We refer to this image as the ‘**all subjects TI**’ image, representing the lower estimate of the predicted TI effect at a particular voxel. We will refer to its voxel values as TI^*^.

**F.7.1** **All-subjects TI image – rationale** **behind computing TI***

We performed finite element analysis using the structural images of each subject to predict the TI magnitude at each voxel. Therefore, for every voxel in the MNI space, we had a predicted TI value from all the subjects. We wanted to create a measure from these that could be compared with the voxel values of (stimTI – shamTI) BOLD fMRI contrast obtained from the second-level analysis, that is, the one-sample t-test across the subjects. One way to do this was to find the mean of the predicted TI values for a voxel across the subjects (TI* = μ). Such a measure, however, ignores the variance in these values. For voxels with comparable mean predicted TI, we would predict high BOLD contrast value for the one that has low variance. The rationale being that for the voxel with high variance, the low predicted TI for some subjects at that voxel location may not necessarily lead to BOLD activation. We also wanted to avoid a measure for which the voxels with very low mean predicted TI but nearly zero variance across the subjects would have high values, which would have been the case with measure of type, TI* = μ/σ. Therefore, we chose TI* = μ - 2*σ as our measure of predicted TI across the subjects that could be compared with the values of the BOLD contrast. Here, μ is mean and σ is the standard deviation of the predicted TI values for a voxel across the subjects. Assuming that the predicted TI values (for a voxel) across the subjects can be approximated by normal distribution, 95.4% of the values would fall within the bounds μ-2*σ and μ+2*σ, and thus this TI* represents the lower estimate of the predicted TI effect at a particular voxel.

**F.8** **Creating mask in frontal lobe for small volume correction of BOLD fMRI activation effects** (Main article – Section 2.2 continued)

This mask was created using the ‘all-subjects TI’ image whose voxel values we refer to as TI*. The TI*was high (>0.6) in the frontal lobe, so we created a mask with MNI y>22 which we used to perform small volume correction of the fMRI results. The values 0.6 is chosen arbitrarily while y>22 was chosen such that it contained the entire region in all-subjects TI image that showed TI^*^>0.6 V/m. The value ‘0.6 V/m’ is high relative to 0.2 V/m which was the average TI^*^ in our target region for which electrode positions were optimized and has a z-score of 4.5, calculated using the mean and standard deviation of the TI^*^ values of all the voxels in all-subjects TI image. Please refer to figure 3B to see the mask and 3C to see the all-subjects TI image after applying this mask.

**F.9** **Creating mask around Cpz electrode location for small volume correction of BOLD fMRI suppression effects** (Main article - section 3.3 continued)

To create the small volume correction mask, the average location of Cpz electrodes across the subjects in MNI coordinates was found which was [1, -57, 90]. This location was outside the brain, as expected. The location in the brain that was closest to this was found which was [0, -57, 78]. The mask was the overlap between the brain (that is, the mask used for the one-sample t-test of StimTI – ShamTI contrast) and a sphere of radius 30 mm around [0, -57, 78]. The rationale behind choosing 30 mm as the radius was that the exact electrode placement location varied across subjects. For a particular subject, the distance of a point having coordinates that are maximally away individually along X, Y, and Z directions from the mean electrode location was found. An average of this was taken across the electrode placements across subjects, which turned out to be 29 mm, and hence a mask of 30 mm radius was formed for small volume correction (Figure 4).
